# Cognitive modelling reveals distinct electrophysiological markers of decision confidence and error monitoring

**DOI:** 10.1101/860379

**Authors:** Manuel Rausch, Michael Zehetleitner, Marco Steinhauser, Martin E. Maier

## Abstract

Is confidence in perceptual decisions generated by the same brain processes as decision itself, or does confidence require metacognitive processes following up on the decision? In a masked orientation task with varying stimulus-onset-asynchrony, we used EEG and cognitive modelling to trace the timing of the neural correlates of confidence. Confidence as reported by observers increased with stimulus-onset-asynchrony in correct and to a lesser degree in incorrect trials, a pattern incompatible with established models of confidence. Electrophysiological activity in two different time periods was associated with confidence, namely, 350 – 500 ms after stimulus onset and 250 – 350 ms after the response. Cognitive modelling revealed that only the activity following on the stimulus exhibited the same statistical regularities as confidence, while the statistical pattern of the activity following the response was not compatible with confidence. It is argued that electrophysiological markers of decision confidence and error awareness are at least in parts distinct.

## Introduction

Decision confidence is a ubiquitous feature of human decision making: Whenever we make a choice, the decision is accompanied by a greater or smaller degree of confidence that the choice is correct. Confidence can be defined as an evaluation of one’s decision making, resulting in a degree of certainty that the decision is correct (Pouget, Drugowitsch, & Kepecs, 2016). How does the brain give rise to confidence? Two conflicting views have been proposed: According to one view, confidence may be generated directly by the very same brain processes that are involved in decision formation (Kepecs, Uchida, Zariwala, & Mainen, 2008; Kiani & Shadlen, 2009; Rolls, Grabenhorst, & Deco, 2010). This view is closely related to Bayesian brain theory, which argues that representations about uncertainty are necessary for optimal choices (Knill & Pouget, 2004). According to the second view, confidence is generated by a separate, metacognitive process that gives rise to both confidence and error awareness (Boldt & Yeung, 2015; Charles & Yeung, 2018). A common mechanism underlying error monitoring and decision confidence may be on-going accumulation of sensory evidence after the decision, allowing observers to reverse their belief about the stimulus (Pleskac & Busemeyer, 2010; Resulaj, Kiani, Wolpert, & Shadlen, 2009; Steinhauser, Maier, & Hübner, 2008; van den Berg et al., 2016).

The aim of the present study was to test if the neural correlates of confidence in a perceptual decision emerge already before the time of the behavioural response consistent with a common origin of confidence and choice formation, or if these correlates do not emerge until the time of neural markers of error awareness following the response. For this purpose, the present study used cognitive modelling and electroencephalography to trace the timing of the neural correlates of confidence in perceptual decisions.

The present study examines three event-related potential (ERP) components that were previously proposed as correlates of confidence: the P3 (Hillyard, Squires, Bauer, & Lindsay, 1971), the error-related negativity ERN (Scheffers & Coles, 2000), and the error-related positivity Pe (Boldt & Yeung, 2015). The P3 is an ERP component recorded over central and parietal electrodes peaking 300 – 500 ms after the presentation of a task-relevant stimulus. It is a natural candidate for a shared electrophysiological correlate of confidence and the decision, because it was suggested that the parietal P3 is a marker of accumulated evidence in perceptual decision making tasks (O’Connell, Dockree, & Kelly, 2012; Philiastides, Heekeren, & Sajda, 2014; Twomey, Murphy, Kelly, & O’Connell, 2015). Previous studies showed that P3 amplitudes are correlated with confidence judgments (Eimer & Mazza, 2005; Hillyard et al., 1971) as well as judgments about subjective visibility (Lamy, Salti, & Barhaim, 2008; Sergent, Baillet, & Dehaene, 2005; Tagliabue et al., 2019). In addition, the P3 showed statistical properties expected from a Bayesian model of decision confidence in a vibrotactile forced-choice task (Herding, Ludwig, von Lautz, Spitzer, & Blankenburg, 2019). However, a marker of accumulated evidence is by far not the only interpretation of the P3: According to a classical theory, the P3 reflects updating of working memory in response to task-relevant events (Donchin & Coles, 1988). Other theories include the global broadcast of visual contents within a neural global workspace (Sergent et al., 2005), the mobilization for action following motivationally significant stimuli (Nieuwenhuis, de Geus, & Aston-Jones, 2011), or a monitoring process if the decision is correctly transformed into an action (Verleger, Ja, & Wascher, 2005)

ERN and Pe are established makers of error processing: If one shared neurocognitive mechanism gives rise to both confidence and error monitoring, confidence should be associated with ERN and Pe. The ERN is an ERP component with frontocentral topography at the same time of shortly after incorrect responses (Falkenstein, Hohnsbein, Hoormann, & Blanke, 1991; Gehring, Goss, Coles, Meyer, & Donchin, 1993). An equivalent yet smaller negativity referred to as CRN was observed after correct responses (Vidal, Burle, Bonnet, Grapperon, & Hasbroucq, 2003). Previous studies suggested that the ERN was associated with participants’ confidence judgments in a flanker task (Scheffers & Coles, 2000). However, the ERN failed to predict graded confidence judgments on a trial-to-trial basis in visual discrimination tasks with briefly flashed stimuli (Boldt & Yeung, 2015). Finally, the ERN can be dissociated from decision confidence by the relation with subjective visibility: In a masked number discrimination task, the ERN varied in an all-or-nothing way and was only present if there was a conscious percept of the stimulus, while confidence varied continuously and did not depend on a conscious percept of the stimulus (Charles, King, & Dehaene, 2014; Charles, Opstal, Marti, & Dehaene, 2013).

The Pe is a parietally focused positive deflection 200 – 500 ms after incorrect responses. The Pe is similar to the parietal P3 in terms of topography and latency despite the fact that the Pe is locked to the response, and P3 to the stimulus (Overbeek, Nieuwenhuis, & Ridderinkhof, 2005). The Pe is a marker of conscious awareness of having committed an error (Nieuwenhuis, Ridderinkhof, Blom, Band, & Kok, 2001) and can be dissociated from the ERN: In a study where participants responded to a masked target stimulus surrounded by visible flanker stimuli, erroneous responding to the flanker elicited only a Pe, but not an ERN (Di Gregorio, Maier, & Steinhauser, 2018). The Pe can be explained by the strength of accumulated evidence of having made an error (Steinhauser & Yeung, 2010, 2012; Ullsperger, Harsay, Wessel, & Ridderinkhof, 2010; Wessel, Danielmeier, & Ullsperger, 2011). Moreover, in a visual discrimination task, the Pe was associated with both confidence in correct responses as well as the subjective belief of having made an error in a gradual way (Boldt & Yeung, 2015). However, the timings of ERN and Pe are not immediately plausible for correlates of decision confidence. As it seems that confidence is experienced already at a point in time when no response has yet been made, correlates of confidence may naïvely be expected before the response, at the same time as the decision or shortly afterwards. And yet, ERN and Pe do not occur until after the response.

How can neural correlates of confidence be identified? If specific neural activity is a correlate of confidence, it must be associated with the same statistical regularities as confidence judgments (Kepecs et al., 2008; Sanders, Hangya, & Kepecs, 2016): By implication, if the statistical regularities of a specific ERP component are incompatible with those of confidence, that component is not a correlate of confidence. In the present study, we tracked the statistical regularities of confidence by fitting a series of cognitive models to the behavioural data. The model that fitted the behaviour best was used to predict the neuronal data. Previous studies used the so-called folded X-pattern as a statistical marker of confidence (Braun, Urai, & Donner, 2018; Fetsch, Kiani, Newsome, & Shadlen, 2014; Herding et al., 2019; Lak, Nomoto, Keramati, Sakagami, & Kepecs, 2017; Urai, Braun, & Donner, 2017). The folded X-pattern is characterised by an increase of confidence with stimulus strength in correct trials and a decrease of confidence with stimulus strength in incorrect trials and was derived from Bayesian decision theory (Hangya, Sanders, & Kepecs, 2016; Sanders et al., 2016), but also follows from signal detection theory (Kepecs et al., 2008) or postdecisional accumulation models (Moran, Teodorescu, & Usher, 2015). However, the folded X-pattern can be misleading about confidence because Bayesian decision theory is compatible with other statistical patterns, too (Adler & Ma, 2018; Rausch & Zehetleitner, 2019b). In addition, in some tasks, confidence empirically increased with stimulus strength in correct trials and to a lesser degree in incorrect trials (Kiani, Corthell, & Shadlen, 2014; Rausch, Hellmann, & Zehetleitner, 2018; Stolyarova et al., 2019; van den Berg et al., 2016), a pattern we refer to as double increase pattern. The double increase pattern can be reproduced by a smaller number of mathematical models, including the weighted evidence and visibility (WEV) model (Rausch et al., 2018), the heuristic detection model (Maniscalco, Peters, & Lau, 2016; Peters et al., 2017), and some Bayesian models (Adler & Ma, 2018; Rausch & Zehetleitner, 2019b). For these reasons, it is not legitimate to assume a specific statistical pattern a priori. However, irrespective of whether confidence follows the folded-X or double increase pattern in a specific task, a neural correlate of confidence should always show the same pattern as the one observed with confidence judgments. In addition, a cognitive model fitted to confidence judgments should also be able to accurately predict the neural correlate of confidence.

To assess the timing of the neural correlates of confidence in perceptual decisions, human observers performed a masked orientation discrimination task (see Fig. 1) while EEG was recorded. After each single response, observers reported their confidence on a scale with the categories, “not at all”, “a little”, “nearly sure”, and “completely sure”. In case observers were aware of an incorrect response, observers were instructed to respond, “not at all”. We used a task where confidence followed the double increase pattern in previous studies (Rausch et al., 2018), because the double increase pattern can be explained by a smaller number of cognitive models. The strength of stimulation was manipulated by varying the stimulus-onset-asynchrony (SOA), i.e. the time between onset of the stimulus and the mask. Bayes factors were used for statistical inference, allowing us to quantify both the evidence for an effect as well as evidence against an effect (Rouder, Speckman, Son, & Morey, 2009).

**Fig. 1.**
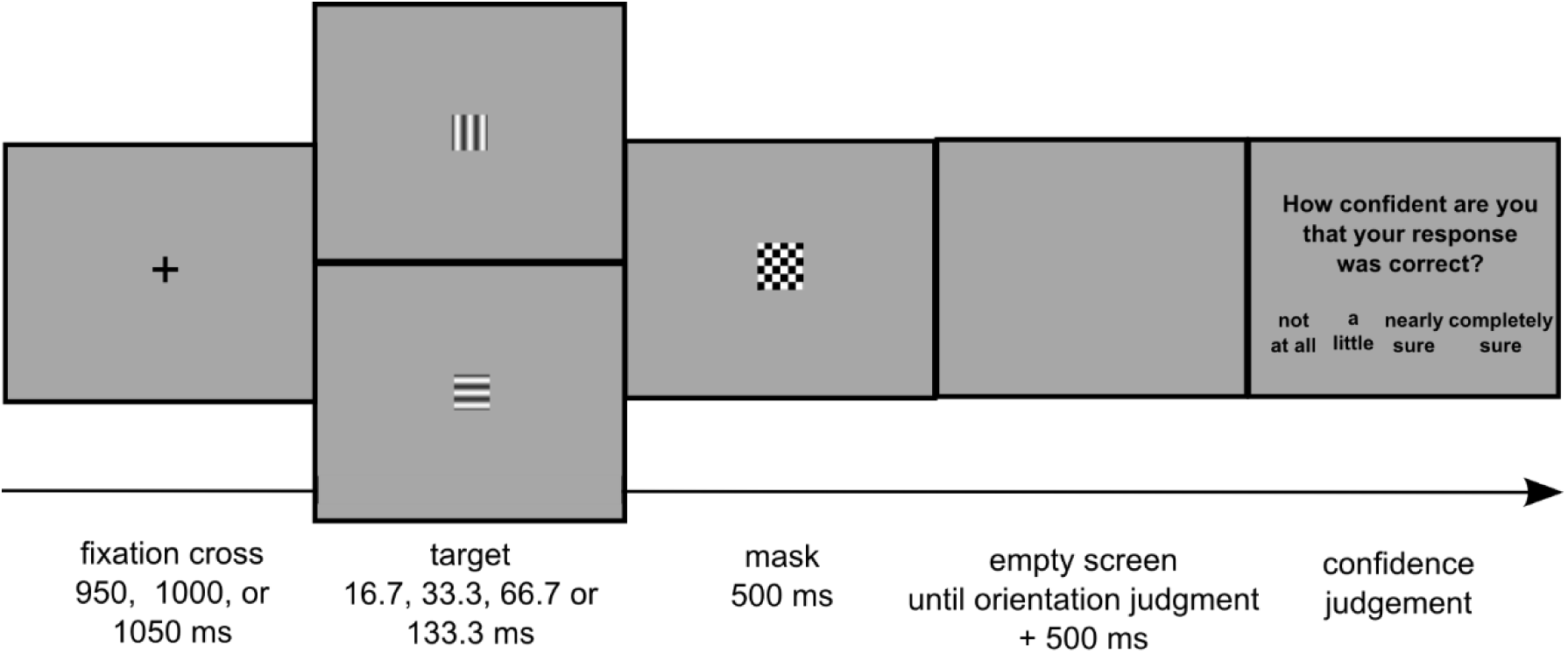
Sequence of events during the Experiment. The target stimulus was a sinusoidal grating, oriented horizontally or vertically. After 16.7, 33.3, 66.7, 133.3 ms, the target was replaced by a chequered mask presented for 500 ms. Afterwards, observers reported first the orientation of the target and then their degree of confidence in having made the correct orientation response. Observers were instructed that accuracy but not speed was critical for both responses.

To trace the statistical regularities underlying confidence, we fitted nine previously proposed models of decision confidence to confidence judgements, including

– SDT (Green & Swets, 1966; Macmillan & Creelman, 2005; Wickens, 2002)
– SDT with noise superimposed on ratings (Maniscalco & Lau, 2016)
– SDT model with postdecisional accumulation (Barrett, Dienes, & Seth, 2013)
– the two-dimensional Bayesian model (Aitchison, Bang, Bahrami, & Latham, 2015)
– the noisy decay model (Maniscalco & Lau, 2016)
– the detection heuristic model (Peters et al., 2017)
– the two high-threshold model (Kellen & Klauer, 2015)
– the two channel model (Rausch & Zehetleitner, 2017)
– the WEV-model (Rausch et al., 2018).

Because the objective of the present study was specifically confidence, we selected models from the literature that are fitted directly to confidence judgments, and not to reaction times. From the model parameters that fitted the behavioural data best, we generated a prediction about the ERP amplitudes under the assumption that the ERP amplitude is proportional to confidence.

With respect to confidence judgments, we expected that confidence increases as a function of the SOA both in correct as well as in incorrect trials, i.e. confidence is characterised by the double increase pattern. Regarding proposed ERP correlates of confidence, as correct responses are commonly associated with more positive activity at the time of the P3 (Koivisto & Revonsuo, 2010), we hypothesized that confidence is positively associated with EEG activity at the time of the P3. As errors are known to cause negative shifts at the time of the ERN, again a positive association was expected between confidence and activity at the time of the ERN (Scheffers & Coles, 2000). In contrast, as errors are known to cause positive shifts at the time of the Pe, we predicted a negative association between confidence and activity at the time of the Pe in line with previous research (Boldt & Yeung, 2015). Moreover, if P3, ERN, and Pe were indeed correlates of confidence, the statistical pattern as a function of SOA and choice accuracy should correspond to the statistical pattern observed in confidence judgments: This means that P3, ERN, and Pe should be characterised by the double increase pattern as well. Regarding cognitive models, we expected that the best fit to the behavioural data should be achieved by one of the models that is in principle able to accommodate the double increase pattern, i.e. the WEV-model, the heuristic detection model, or the noisy decay model. Finally, the models that provide an adequate fit to the behavioural data should also accurately predict the ERP correlates of confidence.

## Methods

### Participants

25 human participants (21 female, 4 male) took part in the experiment. The age of the participants ranged between 18 and 36 years (*Md* = 22). All participants reported normal or corrected-to-normal vision, no history of neuropsychological or psychiatric disorders and not to be on psycho-active medication. All participants gave written informed consent and received either course credits or €8 per hour for participation. The experimental protocol was approved by the ethics committee of the Catholic University of Eichstätt-Ingolstadt.

### Apparatus and stimuli

The experiment was performed a sound-attenuated and electrically shielded cabin. The stimuli were presented on an Iiyama MS103DT monitor with screen diagonal of 51 cm, set at a resolution of 1280 x 1024 px and refresh rate of 60 Hz. The viewing distance, not enforced by constraints, was approximately 60 cm. The experiment was conducted using PsychoPy v.1.83.04 (Peirce, 2007, 2009) on a Fujitsu Celsius W530 desktop computer with Windows 8.1. The target stimulus was a square (size 3° × 3°), textured with a sinusoidal grating with one cycle per degree of visual angle (maximal luminance: 44 cd/m^2^; minimal luminance: 14 cd/m^2^). The mask consisted of a square (4° × 4°) with a black (0 cd/m^2^) and white (60 cd/m^2^) chequered pattern consisting of 5 columns and rows. All stimuli were presented at fixation in front of a grey (29 cd/m^2^) background. The orientation of the grating varied randomly between horizontal or vertical. Participants reported the orientation of the grating with their right hands by pressing the down key when the grating was vertical and the right key when the grating was horizontal. Likewise, participants reported their confidence in being correct with their left hands by pressing one, two, three, or four on the number keys in top row of the keyboard.

### Experimental trial

Each trial began with the presentation of a fixation cross whose duration randomly chosen between 950, 1000, and 1050 ms. Then the target stimulus was shown for a short period of time until it was replaced by the masking stimulus. There were four different possible SOAs, time periods between target onset and mask onset, 16.7, 33.3, 66.7 and 133.3 ms. The mask was presented for 500 ms. When the mask had disappeared, an empty screen was shown. Participants then indicated whether the target had been horizontal or vertical. The question “How confident are you about your response?” was displayed on screen 500 ms after the response, with the four response options “not at all”, “a little”, “nearly sure”, and “completely sure”. Participants then pressed a key to indicate their degree of confidence that their orientation response was correct. In case of incorrect orientation responses, the trial ended by the presentation of the word *error* for 1,000 ms.

### Design and procedure

Participants were instructed to report the orientation of the grating as accurately as possible without time pressure and to guess the orientation of the target if they had no idea about the orientation at all. In addition, they were instructed that they should report their degree of confidence that their orientation response had been correct, they should report their confidence as accurately as possible and that if they were aware that they had made an error, they should rate their degree of confidence as “not at all”.

The experiment consisted of one training block and 24 experimental blocks of 40 trials each. Each SOA featured 10 times in each block in random order. The orientation of the target stimulus varied randomly across trials. After each block, the percentage of errors was displayed to provide participants with feedback about their accuracy.

### EEG acquisition

The electroencephalogram (EEG) was recorded from 64 electrodes using a BIOSEMI Active-Two system (BioSemi, Amsterdam, The Netherlands; Ag/AgCl electrodes, channels Fp1, AF7, AF3, F1, F3, F5, F7, FT7, FC5, FC3, FC1, C1, C3, C5, T7, TP7, CP5, CP3, CP1, P1, P3, P5, P7, P9, PO7, PO3, O1, Iz, Oz, POz, Pz, CPz, Fpz, Fp2, AF8, AF4, AFz, Fz, F2, F4, F6, F8, FT8, FC6, FC4, FC2, FCz, Cz, C2, C4, C6, T8, TP8, CP6, CP4, CP2, P2, P4, P6, P8, P10, PO8, PO4, O2 as well as the left and right mastoid, relative to common mode sense CMS active electrode and driven right leg DR passive electrode). Vertical and horizontal electrooculogram (EOG) was recorded from electrodes above and below the right eye and on the outer canthi of both eyes. All electrodes were off-line re-referenced to linked mastoids. EEG and EOG data were continuously recorded at a sampling rate of 512 Hz.

### EEG analysis

The analysis of the EEG data was performed using MNE-Python (Gramfort et al., 2013, 2014). First, the data was re-referenced to the linked mastoids. Next, the signal was band-pass filtered between 0.5 and 40 Hz by windowed finite impulse response filtering. The raw data was inspected visually to remove extreme noise events and artefact-contaminated electrodes. Then, we conducted an independent component analysis based on the fastica algorithm (Hyvärinen, 1999), identifying components representing blinks and/or horizontal eye movements and removing these artefacts before back-projection of the residual components.

For analysis of the P3 time window, the continuous EEG was epoched into segments starting 200 ms before stimulus onset and ending 600 ms after stimulus onset. The 200 ms pre-stimulus interval was used for baseline correction. For analyses of the ERN and Pe time windows, the EEG data was epoched into segments starting 200 ms before the orientation response and ending 600 ms after the response. The time range between 150 and 50 ms before the response was used as baseline correction. Epochs with amplitude changes greater than 100 μV were excluded from analysis, the same exclusion criterion as in a previous study of EEG correlates of confidence (Boldt & Yeung, 2015). Finally, ERP waveforms were obtained by averaging across epochs (but not for the validation of the prediction by the cognitive models, see below). EEG activity in specific time windows was quantified by calculating mean amplitudes because mean amplitudes are robust to different numbers of trials across conditions (Luck, 2014). The time windows were 350 – 500 ms poststimulus at electrode Pz for the P3, −40 – 60 ms after the orientation discrimination response at electrode FCz for the ERN, and 250 – 350 ms after the orientation discrimination response at electrode Pz for the Pe, the same time windows as in a previous study of EEG correlates of confidence (Boldt & Yeung, 2015). The time window of 350 – 500 after stimulus onset excluded the point in time when participants responded to the orientation of the stimulus (reaction time at the SOA of 133.3 ms: *M* = 624.6 ms, at the SOA of 16.7 ms: *M* = 1001.0 ms). As can be seen from Fig 5L, the Pe in the present study seemed to stretch over a more prolonged time window than 250 – 350 ms postresponse. For this reason, we repeated all analyses using a time window of 200 – 500 ms postresponse, which came to the same interpretation of the data. After artefact rejection, there were on average *M* = 816.3 stimulus-locked epochs and *M* = 800.0 response-locked epochs per participant. In the condition with the smallest number of trials (incorrect trials at the SOA of 133.3 ms), there were on average 10.1 trials. To create topographical maps, artefact-contaminated electrodes that were excluded in the beginning were interpolated using spherical splines (Perrin, Pernier, Bertrand, & Echallier, 1989).

**Fig. 2.**
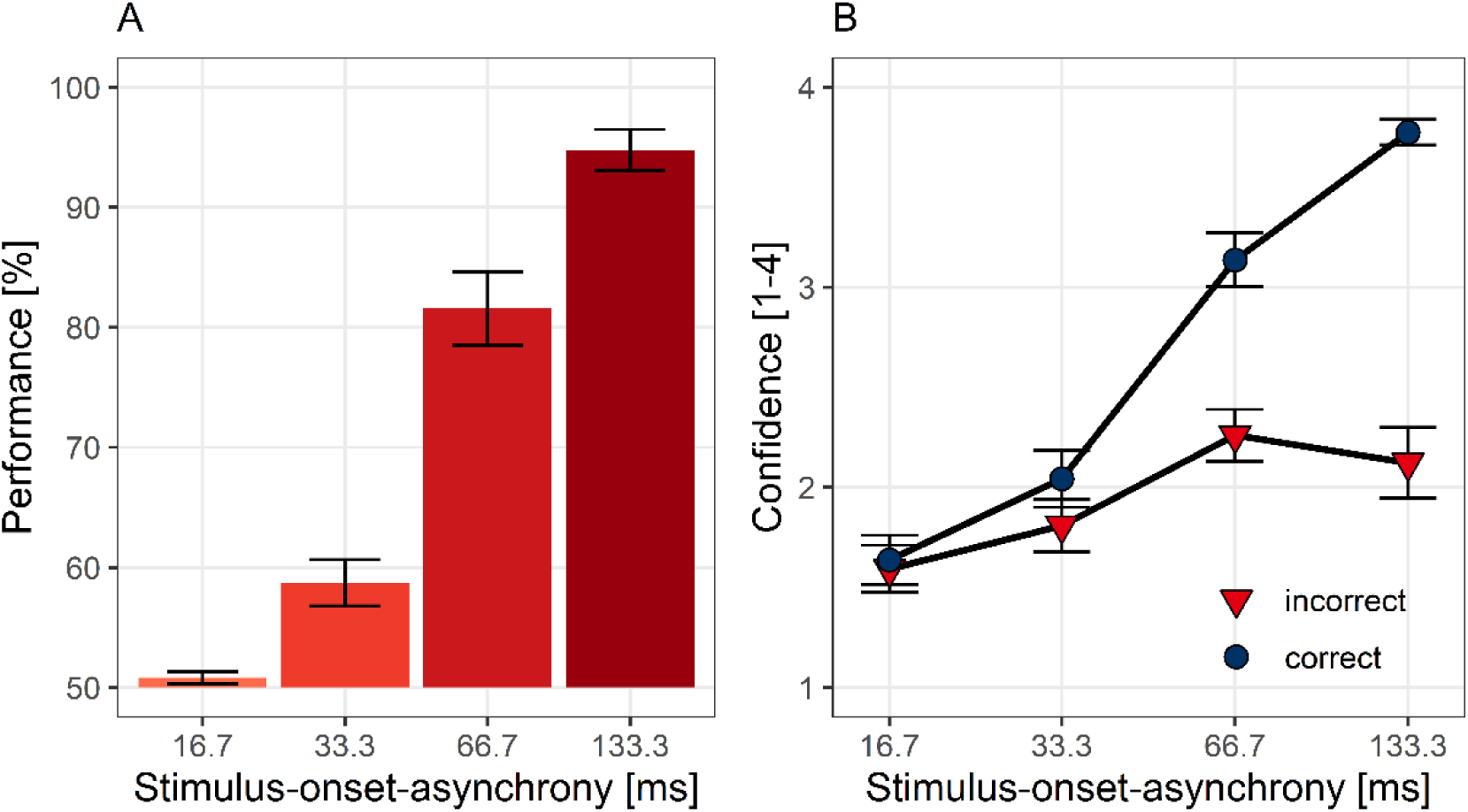
(A) Accuracy in the orientation discrimination task depending on stimulus-onset-asynchrony. B): Decision confidence as a function of stimulus-onset-asynchrony in correct (blue symbols) and incorrect trials (red). Bars and symbols indicate observed means. Error bars indicate 1 *SEM*.

**Fig. 3.**
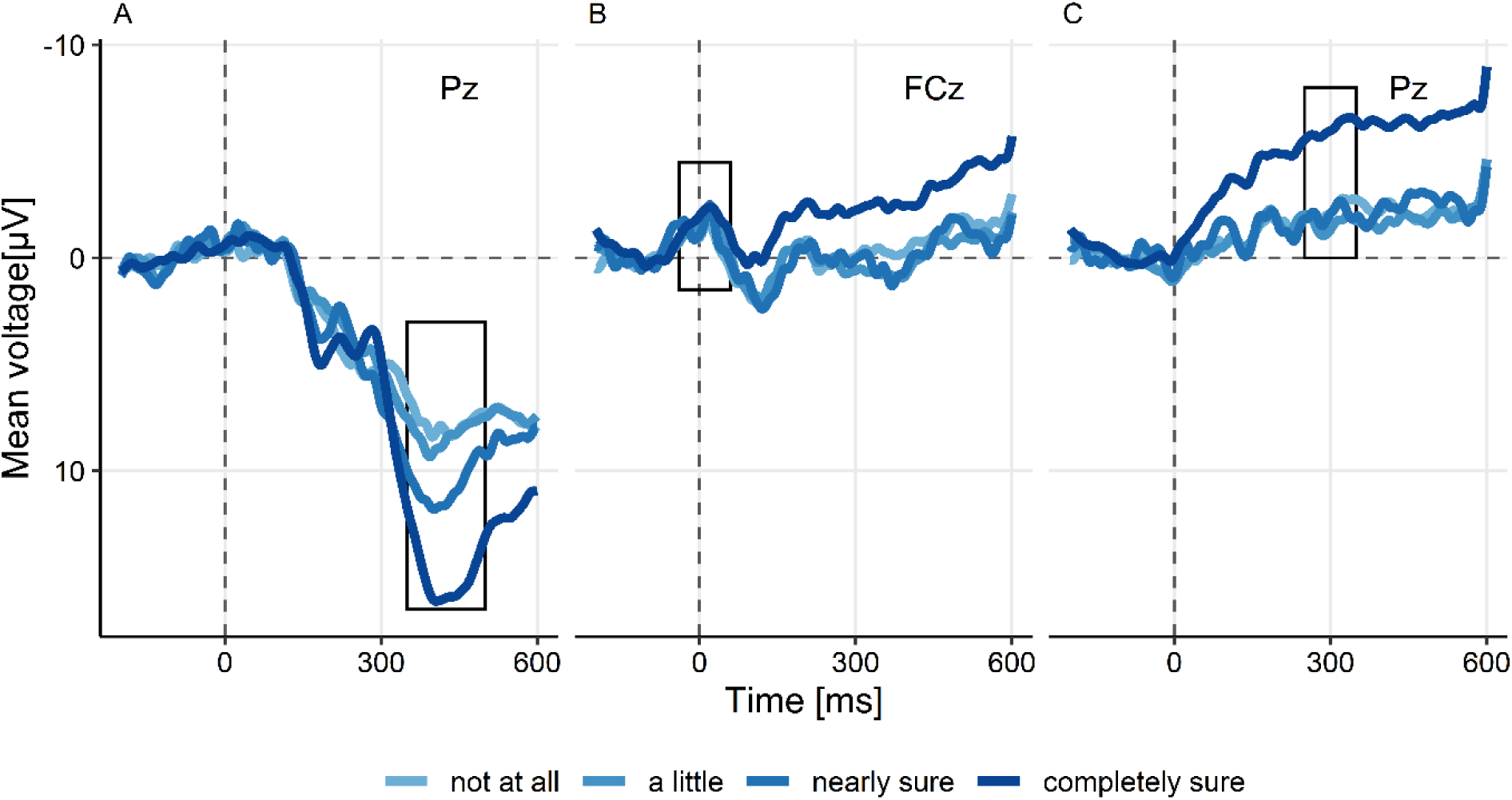
Event-related potentials in correct trials as a function of confidence judgments. (A) Activity at the parietal electrode Pz locked to the onset of the target stimulus. The box highlights the range of the P3 time window. The ERP is locked to the onset of the target stimulus. (B) Activity at the frontocentral electrode FCz locked to the orientation response. The box highlights the time range of the ERN. (C) Activity at Pz locked to the orientation response. The box highlights the time range of the Pe. Colours indicate the degree of confidence reported by the observers; the darker the blue, the more confident the observers were.

**Fig. 4.**
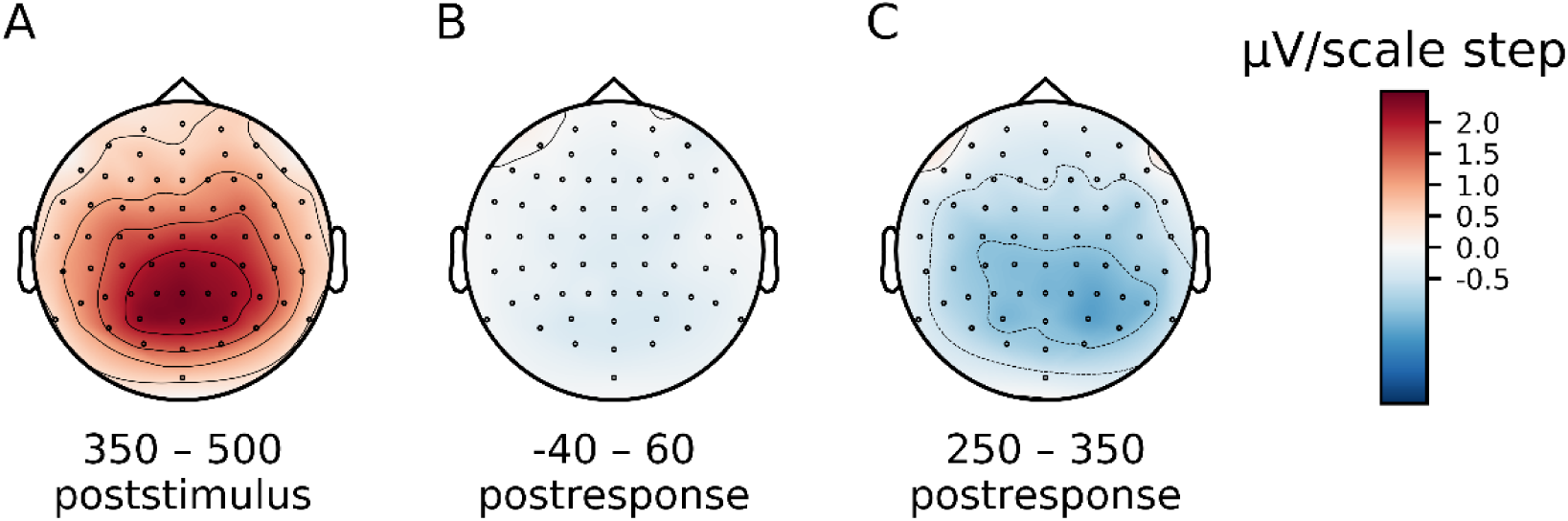
Distribution of the association between ERPs and confidence in correct trials across the scalp. Maps are based on regression slopes with ERP amplitudes as a function of confidence (A) Time window 350 – 500 ms after target stimulus onset. (B) Time window between 40 ms before the orientation response and 60 ms after the response. (C) Time window 250 – 350 ms after the response.

**Fig. 5.**
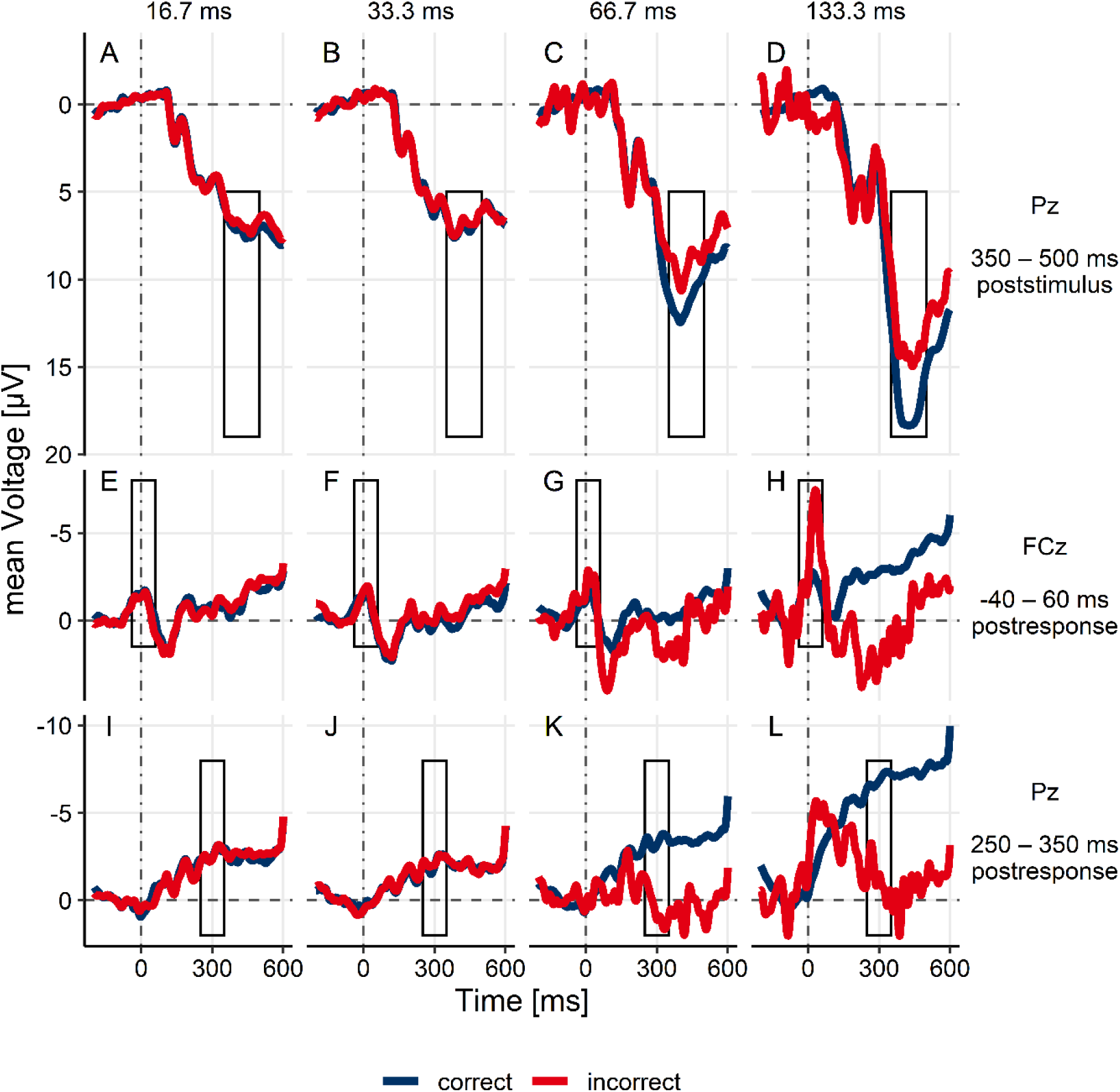
Event-related potentials as a function of stimulus-onset-asynchrony (different columns) and accuracy of the orientation response (blue: correct responses; red: incorrect responses). (A-D) ERP activity locked to target stimulus onset, recorded at Pz. The box highlights the time window of the P3 (350 – 500 ms poststimulus). (E-H) ERP activity locked to the orientation response, recorded at FCz. The box highlights the time window of the ERN (−40 – 60 ms postresponse). (I-L) ERP activity locked to the orientation response, recorded at Pz. The box highlights the time window of the Pe (250 – 350 ms postresponse).

### Model specification

Nine models were fitted to the combined distributions of orientation discrimination and confidence judgments, separately for each single participant.

i. SDT
ii. SDT with noise superimposed on ratings
iii. Noisy decay model
iv. WEV-model
v. Two channel model
vi. SDT model with postdecisional accumulation
vii. Detection heuristic model
viii. Two-dimensional Bayesian model
ix. Two high-threshold model

For all nine models, we assumed that the stimulation was comprised of two experimental variables, the identity of the stimulus *S*_*id*_ and the strength of the stimulus *S*_*s*_. Participant were assumed to select a discrimination response *R*_*id*_ ∈ {0,1} about the identity of the stimulus *S*_*id*_ ∈ {0,1}. Models (i)-(vi) were derived from SDT and assumed the same architecture for the choice about the identity of the stimulus. In contrast, models (vii)-(ix) were based on different decision architectures.

#### SDT derived models

Models (i) -(vi) assumed that a decision about the identity of the stimulus was made based on a comparison between a continuous decision variable for the discrimination judgment *δ*_*id*_ with the free criterion parameter *θ*_*id*_. Participants responded *R*_*id*_ = 0, when *δ*_*id*_ < *θ*_*id*_, and *R*_*id*_ = 1 if *δ*_*id*_ > *θ*_*id*_. The decision variable for the discrimination judgment *δ*_*id*_ was modelled as a random sample from a Gaussian distribution 𝒩:

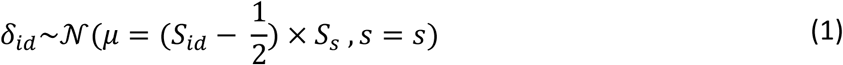

The stimulus strength *S*_*s*_ was a free parameter specific to each SOA. When *S*_*id*_ = 0, the distribution of *δ*_*id*_ was shifted to the left by the distance of *S*_*s*_/2. When *S*_*id*_ = 1, the distribution is shifted by the same distance to the right. Thus, *S*_*s*_ denotes the distance of the distributions generated by the two possible identities of the stimulus and is in this respect equivalent to the sensitivity parameter in standard SDT. Concerning the standard deviation *s*, model fitting was reproduced with two different assumptions about the variability of *δ*_*id*_: For the first set of analyses, the standard deviation of *δ*_*id*_ was fixed at 1 for both identities of the stimulus. For the second set of analyses, the variability of *δ*_*id*_ could vary depending on *S*_*id*_: An additional parameter *λ* indicated the ratio of the standard deviations of *δ*_*id*_ created by two possible identities of the stimulus.

A specific degree of confidence *C* ∈ {1, 2, 3, 4} was determined by comparing the decision variable for confidence *δ*_*c*_ against a set of three criteria *θ*_*c*_. Each criterion delineated between two adjacent categories of confidence, e.g. participants selected the category 2 if *δ*_*c*_ fell between *θ*_*c*1_ (which separated category 1 and 2) and *θ*_*c*2_ (which separated category 2 and 3). To be consistent with standard SDT, we fitted three different criteria for each of the two response options. The different models were characterised by different ways how *δ*_*c*_ was determined.

##### SDT rating model

According to model (i), the decision variables for identification and confidence were identical:

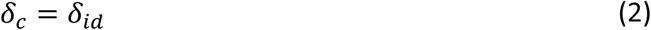

##### Noisy SDT model

According to model (ii), *δ*_*c*_ was sampled from a Gaussian distribution, with a mean equal to the decision variable *δ*_*id*_ and the standard deviation *σ*, which was an additional free parameter:

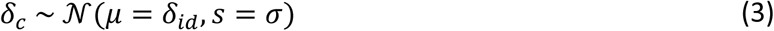

##### Noisy decay model

According to model (iii), the, *δ*_*c*_ was also sampled from a Gaussian distribution with the standard deviation *σ*. The mean of *δ*_*c*_ depended on *δ*_*id*_, but *δ*_*id*_ was reduced by multiplication with a signal reduction parameter *ρ*_*S*_. The signal reduction parameter *ρ*_*S*_ was a separate free parameter for each SOA and was bounded between 0 and 1.

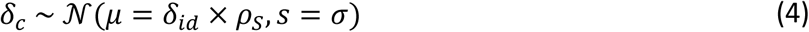

##### WEV model

According to models (iv), *δ*_*c*_ was again sampled from a Gaussian distribution with the standard deviation *σ*:

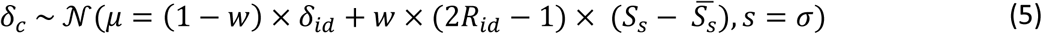

Formula (5) ensured that the centre of the distribution was shifted towards 0 when the strength of stimulation *S*_*s*_ was low and away from 0 when *S*_*s*_ was high. The parameter *w* captured the degree to which participants relied on sensory evidence about the identity or on identity-irrelevant evidence when they determined their degree of confidence. When *w* = 0, the model was identical to the noisy SDT model; when *w* = 1, *δ*_*c*_ only depended on the strength of stimulation *S*_*s*_, but not on the decision variable for the identification judgment *δ*_*id*_. The term 2*R*_*id*_ − 1 ensured that strong stimuli tended to shift the location of the distribution in a way that high confidence was more likely, and likewise, weak stimuli tended to shift the location of the distribution in a way that the probability of low confidence increased. 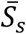 denotes the mean of *S*_*s*_ across the five SOAs and was added to the formulae to increase stability during parameter fitting.

##### Two-channel model

According to model (v), *δ*_*c*_ was again sampled from a Gaussian distribution, but now independently from *δ*_*id*_:

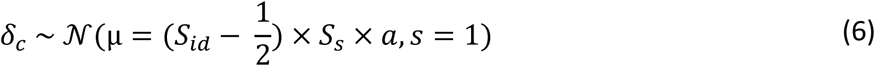

The free parameter *a* expressed the fraction of signal available to the second channel relative to the signal available to the first channel.

##### SDT model with postdecisional evidence

According to model (vi), the, *δ*_*c*_ was again sampled from a Gaussian distribution:

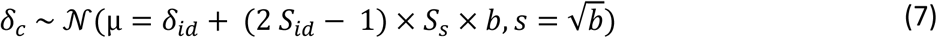

The free parameter *b* indicated the amount of postdecisional accumulation, and the term 2 *S*_*id*_ − 1 ensured that postdecisional accumulation tended to decrease *δ*_*c*_ when *S*_*id*_ = 0, and to increase *δ*_*c*_ when *S*_*id*_ = 1.

#### Non-SDT models

Model (vii)-(ix) assumed a different decision architecture for the identification judgment than models (i)-(vi).

##### Detection heuristic model

According to model (vii), there were two separate decision variables for the identification judgment, each belonging to one possible identity of the stimulus:

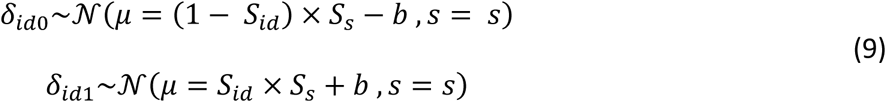

The parameter *b* reflected the a priori bias in favour of *R*_*id*_ = 1. Participants were assumed to respond *R*_*id*_ = 0, when *δ*_*id*0_ > *δ*_*id*1_, and *R*_*id*_ = 1 if *δ*_*id*0_ < *δ*_*id*1_. Confidence judgments were only based on the decision variable pertaining to the selected response: When *R*_*id*_ = 0, *δ*_*id*0_ was compared against a series of confidence criteria *θ*_*c*0_ to select a specific degree of confidence; and when *R*_*id*_ = 1, the comparison was based on *δ*_*id*1_ as well as a second set of criteria *θ*_*c*1_.

##### 2-D Bayesian model

According to model (viii), there were again two separate decision variables, *δ*_*id*0_ and *δ*_*id*1_, referred to as ‘sensory signals’ by Aitchinson et al. (2015), each referring to one of the two possible identities of the stimulus:

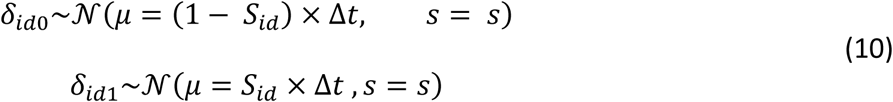

*Δt* denotes the physical SOA and s is a free noise parameter. The model assumed that the observer’s choices about the identity of the stimulus and about the visibility depended on the posterior probability of the identity of the stimulus given the decision variables

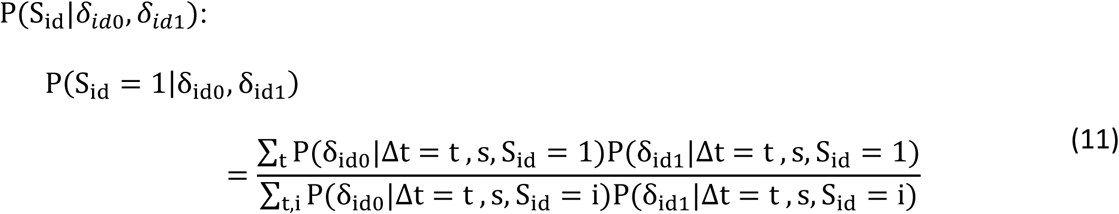

A specific identity and degree of visibility were chosen by comparing the posterior probability P(S_id_ = 1|*δ*_*id*0_, *δ*_*id*1_) against a set of criteria *θ*. It was assumed that the possible identities and degrees of visibility formed an ordered set of decision options. Each criterion delineated two adjacent decision options, e.g. participants chose to respond that the identity was 1 and visibility was 1 if P(S_id_ = 1|δ_id0_, δ_id1_) was smaller than the criterion associated with identity 1 and visibility 2, and at the same time P(S_id_ = 1|δ_id0_, δ_id1_) was greater than the criterion for identity 0 and visibility 1. Finally, it was assumed that observers did not always give the same response as they intended to. When a lapse occurred, identification and visibility responses were assumed to be random with equal probabilities. The lapse rate λ was an additional free parameter.

##### Two high thresholds model

Model (ix), the two high thresholds model, assumed that the decision variable for the identification judgment *δ*_*id*_ was not continuous, but categorical *δ*_*id*_ ∈ {0, 0.5, 1}: Observer could either detect the identity of the stimulus and choose the response accordingly *R*_*id*_ = 0 *if δ*_*id*_ = 0, and *R*_*id*_ = 1 *if δ*_*id*_ = 1. Alternatively, observers could be in a state of uncertainty, *δ*_*id*_ = 0.5, in which no information about the identity was available, and observers responded by random guessing. The probability to detect the identity of the stimulus depended on the five SOAs as well as on the identity of the stimulus, resulting in a total of ten detection parameters *p*(*δ*_*id*_ = *S*_*id*_|*S*_*s*_, *S*_*id*_). A guessing parameter *g* determined the probability with which observers responded *R*_*id*_ = 1 when they were in the state of uncertainty. A specific degree of confidence was sampled randomly depending on the three possible states of *δ*_*id*_ and the response *R*_*id*_. As the response was fixed when observers detected the identity, there were four different sets of probabilities to determine confidence judgments *p*(*C* = *c*|*δ*_*id*_ = 0), *p*(*C* = *c*|*δ*_*id*_ = 1), *p*(*C* = *c*|*δ*_*id*_ = 0.5, *R*_*id*_ = 0), and *p*(*C* = *c*|*δ*_*id*_ = 0.5, *R*_*id*_ = 1). All *p*(*δ*_*id*_ = *S*_*id*_|*S*_*s*_, *S*_*id*_), *p*(*C* = *c*|*δ*_*id*_, *R*_*id*_) and *g* were free parameters.

### Model fitting

The nine models were fitted to the combined distributions of orientation discrimination and confidence judgments separately for each single participant. First, the frequency of each confidence category was counted for each orientation of the stimulus and each orientation response. Then, for each model, the set of parameters was determined that minimized the negative log-likelihood. For models (i)-(vii) and (ix), the likelihood was calculated analytically. Only for the 2-D Bayesian model, the likelihood was approximated by simulation. Minimization was performed using a general SIMPLEX minimization routine (Nelder & Mead, 1965). To quantify the goodness-of fit of the nine models, we calculated the BIC (Schwarz, 1978) and the AICc (Burnham & Anderson, 2002), a variant of the Akaike information criterion (Akaike, 1974) using the negative likelihood of each model fit with respect to each single participant and the trial number.

### Predictions of ERP amplitudes

To determine expected ERP amplitudes from model fits, we first used the parameter sets obtained during model fitting of the behavioural data to calculate the probabilities of all four confidence categories as a function of SOA and choice accuracy. Then, the expected ERP amplitude was determined as the sum of the transformed confidence categories, weighted by the probability of each confidence category. The transformations of confidence were determined by SIMPLEX minimization of the sum of squares of the deviation between the expected ERP amplitude and the observed ERP amplitudes across trials separately for each single subject. Two types of transformations of confidence were used: A linear transformation and a monotonous transformation. For the monotonous transformation, there was one free parameter for each of the four confidence categories, specifying the expected ERP amplitude. The fitting algorithm was constrained to ensure that the expected ERP amplitude was either monotonously increasing or decreasing with confidence. Finally, the correlations between predicted and observed ERP amplitudes were assessed across trials separately for each participant.

### Bayesian statistics

All statistical tests were based on Bayes factors (Rouder et al., 2009), as implemented in the R package *BayesFactor* (Morey & Rouder, 2015). To test if an ERP component was related to confidence or to the SOA, we used a Bayesian linear mixed regression model with a fixed effect of SOA or confidence and a random effect of participant on the intercept, using default mixture-of-variance priors and a scale parameter of r = 1/2 (Rouder & Morey, 2012). Conceptually, the prior represents the a priori belief that smaller regression slopes are more plausible than large slopes, although even very large slopes are not impossible. Each Bayes factor represents a comparison between the full regression model and a regression model with only the random effect of participant. To compare fits between models of confidence, the Bayesian equivalent of a paired t-test was used, assuming a Cauchy distribution with a scale parameter of 1 as prior for the standardized effect size δ, a choice recommended as default (Rouder et al., 2009). The strength of statistical evidence was interpreted according to an established guideline (Lee & Wagenmakers, 2013). In addition, we constructed 95% HDI intervals of the regression slopes or mean differences by 10^6^ samples from the posterior distribution using the same models and priors as for Bayes factors.

## Results

### Behavioural results

Discrimination performance of the orientation ranged between chance at the shortest SOA (*M* = 50.8%, *SD* = 2.6) and close-to-ceiling at the longest SOA (*M* = 94.8%, *SD* = 8.5, see Fig. 2A). Confidence ranged between *M* = 1.6 (*SD* = 0.6) on a four-point scale at the shortest SOA and *M* = 3.7 (*SD* = 0.4) at the longest SOA. Fig. 2B shows that confidence was characterised by an increase with SOA in correct as well as in incorrect trials. The evidence for the increase with SOA was extremely strong for correct trials, 95% HDI [0.016 0.020] scale steps/ms, *BF*_*10*_ = 1.3·10^24^, and strong for incorrect trials, 95% HDI [0.002 0.007] scale points/ms, *BF*_*10*_ = 23.7.

### ERP results

The effects of confidence were examined in correct trials during the time windows of the three candidate correlates of confidence: P3, ERN, and Pe. Consistent with our prediction, there was extremely strong evidence that EEG activity in the P3 time range (350 – 500 ms after onset of the target stimulus, recorded at the parietal electrode Pz) increased with confidence, 95% HDI [1.8 2.9] µV/scale step, *BF*_*10*_ = 3.6·10^10^ (see Fig. 3A). Fig. 4A shows that the association between ERPs and confidence in correct trials during the P3 time window had a centroparietal distribution over the scalp, consistent with known topographies of the P3 in difficult perceptual discrimination tasks (Koivisto & Revonsuo, 2010).

Fig. 3B shows the effect of confidence in correct trials during the ERN time window (−40 – 60 ms after the orientation response, at the frontocentral electrode FCz). The evidence for an effect of confidence was not conclusive, 95% HDI [−0.6 0.1] µV/scale step, *BF*_*10*_ = 0.45. However, although a positive relation between confidence and ERN would have been expected, the trend was in the opposite direction. As shown by Fig. 4B, no clear effect of confidence emerged anywhere over the scalp at the time of the ERN. Finally, Fig. 3C shows the effect of confidence in correct trials during the Pe time window (250 – 350 ms after the orientation response, at Pz). In line with our prediction, there was strong evidence that confidence was negatively associated with the ERP at the time of the Pe, 95% HDI [−1.8 - 0.5] µV/scale step, *BF*_*10*_ = 60.6 µV. As can be seen from Fig. 4C, the association between ERPs and confidence in correct trials during the Pe time window had a posterior parietal distribution over the scalp, consistent with known topographies of the Pe (Boldt & Yeung, 2015).

Next, we tested if P3, ERN and Pe are markers of confidence by comparing the effects of SOA on confidence judgments and on ERPs, separately for correct and incorrect trials. Consistent with the pattern of confidence judgments, there was extremely strong evidence that the ERP in the P3 range increased with SOA in correct trials, 95% HDI [0.08 0.10] µV/ms, *BF*_*10*_ = 1.8·10^25^, as well as in incorrect trials, 95% HDI [0.04 0.08] µV/ms, *BF*_*10*_ = 1.2·10^7^ (see Fig. 5A-D, see also Fig. 7). In the ERN range, Fig. 5H shows that the strongest effect at the time of the ERN was a negative shift in incorrect trials at the longest SOA. The effect of SOA in incorrect trials was in the opposite direction as the pattern of confidence judgments: the evidence was extremely strong for a negative, not positive shift, 95% HDI [−0.04 −0.01] µV/ms, *BF*_*10*_ = 164.5. The evidence with respect to an effect on correct trials was not conclusive, 95% HDI [−0.01 0.00] µV/ms, *BF*_*10*_ = 2.9. Fig 5I-L show that the EEG activities in correct and incorrect trials at the time of the Pe seemed to diverge from each other with increasing SOA, i.e. the Pe seemed to follow the folded X-pattern. This is inconsistent with double increase pattern of confidence judgments, as confidence had increased with SOA in both correct and incorrect trials. As EEG activity at the time of the Pe is negatively associated with confidence, the pattern of confidence judgments implied a negative shift with SOA in both correct and incorrect trials. In accordance with the pattern of confidence judgments, there was extremely strong evidence for the negative shift with increasing SOA in correct trials, 95% HDI [−0.05 −0.03] µV/ms, *BF*_*10*_ = 7.6·10^7^. However, although the pattern of confidence judgments implied a negative shift with increasing SOA in incorrect trials as well, there was moderate evidence against a relationship between SOA and ERPs in incorrect trials at the time of the Pe, 95% HDI [−0.01 0.03] µV/ms, *BF*_*10*_ = 0.29.

**Fig. 6.**
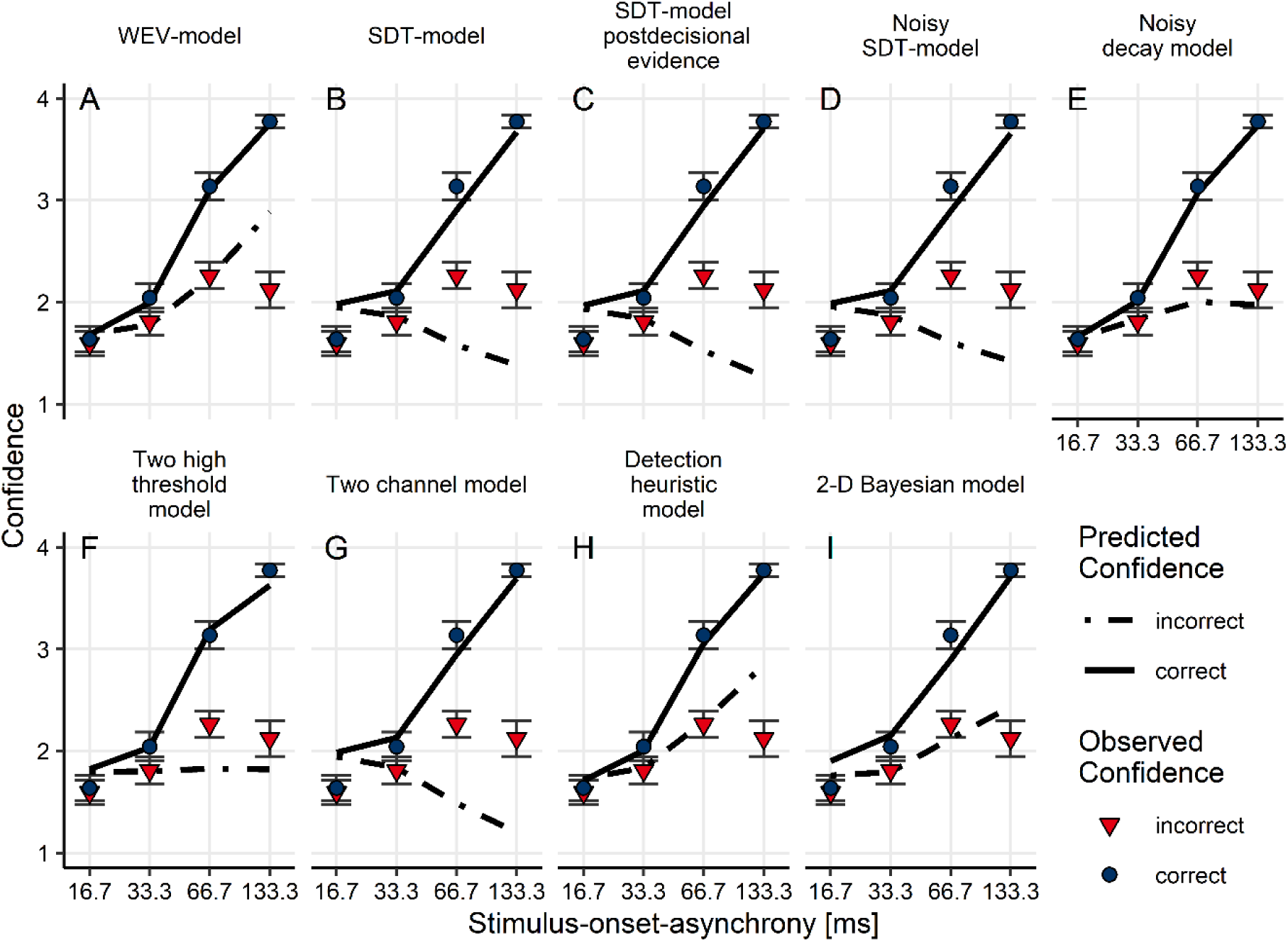
Mean confidence judgments depending on stimulus-onset-asynchrony (x-Axis) and accuracy of the orientation response. Different panels show the prediction of the different models based on the sets of parameters identified during model fitting, assuming constant variances of the decision variable. Solid lines indicate the prediction for correct trials, dashed lines for incorrect trials. Blue circles indicate observed confidence judgments in correct trials, and red triangles in incorrect trials. Error bars = 1 *SEM*.

**Fig. 7.**
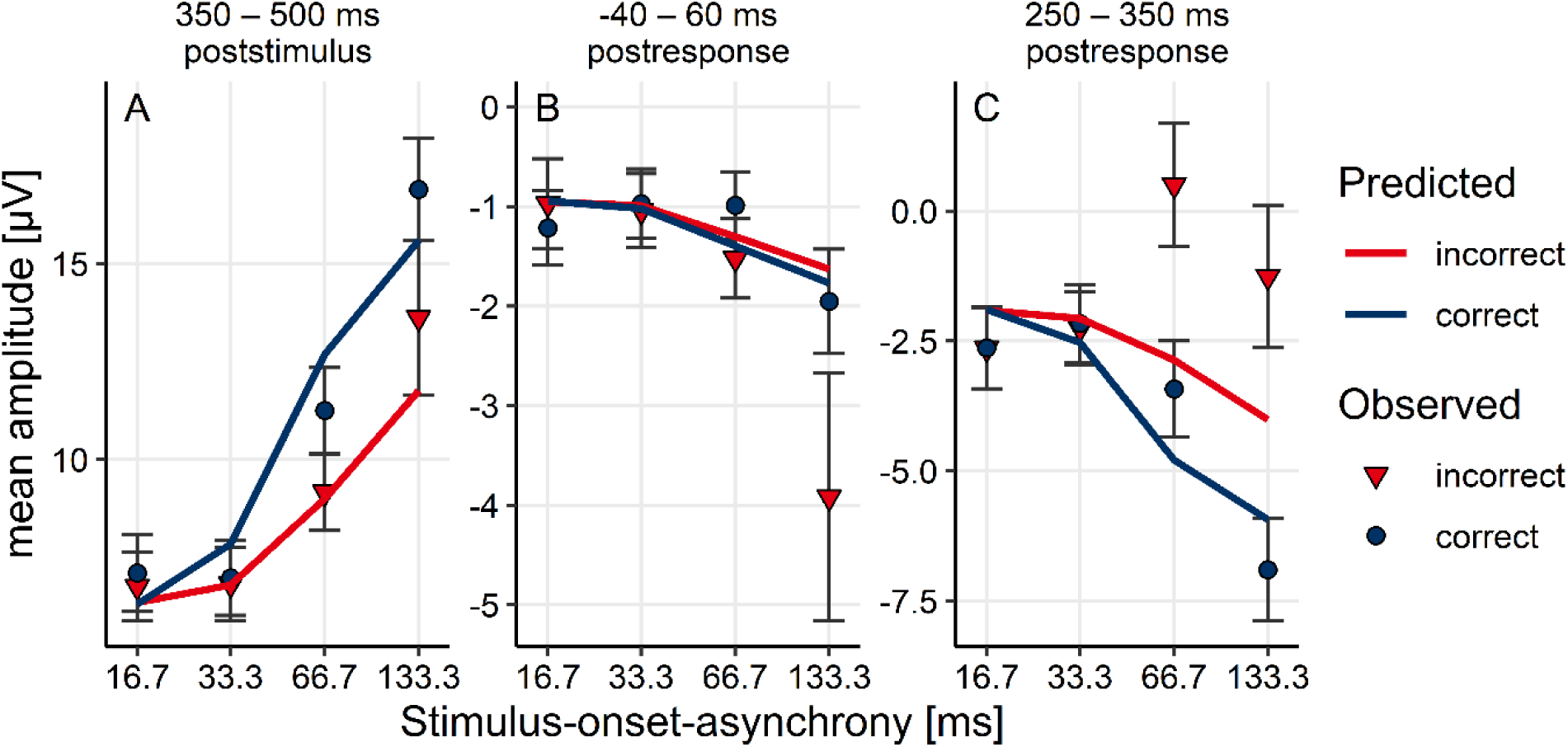
Comparison between predicted and observed amplitudes (A) in the P3 time window, (B) in the ERN time window, and (C) in the Pe time window, depending on stimulus-onset-asynchrony (x-Axis) and accuracy of the orientation response (colours). Symbols indicate observed data. The prediction was determined based on the parameters of the WEV-model fitted to confidence judgments as well as a linear transformation of confidence. Error bars = 1 *SEM*.

### Cognitive modelling

#### Modelling confidence judgments

Fig. 6 shows confidence judgments as a function of SOA and discrimination accuracy compared to the model prediction based on parameter sets identified during fitting. The WEV-model, the noisy decay model, the detection heuristic model and the 2-D Bayesian model correctly predicted that confidence in incorrect trials increases with SOA (Fig. 6, A, E, H, I). The other models could not reproduce the double increase pattern (Fig. 6, B, C, D, F, G).

Quantifying model fit using the Akaike information criterion (AIC_c_) and the Bayes information criterion (BIC) showed that the best fit to the data was obtained by the WEV-model, followed by the noisy decay model (see Table 1). Regarding AIC_c_, the evidence that the WEV model performed better than the noisy decay model was not conclusive, but there was very strong evidence that the WEV-model performed better than the two-high-thresholds-model and extreme evidence that the WEV-model performed better than each of the other models. Regarding BIC, there was moderate evidence that the WEV-model performed better than the detection heuristic model, strong evidence that the WEV-model was better than the noisy decay model, and extreme evidence that the WEV model was better than each of the other five models. These results were essentially the same when it was assumed that the variances of the decision variable differed between horizontal or vertical stimuli. To assess the accuracy of model classification, models were fitted to simulated data based on the parameter sets obtained during model fitting of the empirical data. It was found that model classification was suitable for the purpose of the present study.

**Table 1.** Comparisons of goodness-of-fit of each model against the WEV-model

| Model | Akaike Information criterion |  |  | Bayes information criterion |  |  |
| --- | --- | --- | --- | --- | --- | --- |
| | $BF_{10}^*$ | $\Delta AIC$ | $\Delta AIC$ | $BF_{10}^*$ | $\Delta BIC$ | $\Delta BIC$ |
|  |  | 2.5%** | 97.5%** |  | 2.5%** | 97.5%** |
| SDT model | $1.6 \cdot 10^3$ | -186.8 | -81.0 | 658.3 | -177.2 | -71.6 |
| SDT with postdecisional evidence model | 832.3 | -174.9 | -72.0 | 528.7 | -170.3 | -67.4 |
| Noisy SDT | $1.2 \cdot 10^3$ | -186.8 | -79.0 | 760.9 | -182.1 | -74.5 |
| Noisy decay model | 0.35 | -21.7 | 4.9 | 16.5 | -35.8 | -8.5 |
| Two-high-threshold model | 43.3 | -146.7 | -42.8 | $1.4 \cdot 10^3$ | -185.2 | -79.7 |
| Two-channel model | $1.2 \cdot 10^3$ | -184.5 | -78.4 | 791.1 | -179.8 | -73.7 |
| Detection heuristic model | 101.3 | -49.0 | -16.2 | 6.8 | -39.4 | -7.1 |
| 2D Bayesian model | $7.9 \cdot 10^4$ | -193.9 | -104.1 | $1.1 \cdot 10^4$ | -174.5 | -85.0 |

\* $BF_{10}$ : Bayes factor comparing the hypothesis that the WEV-model and the model in the row are associated with different mean AIC<sub>c</sub> or BIC, respectively, against the hypothesis that the means were the same, assuming a Cauchy prior distribution of the effect size with a scale parameter of 1 (Rouder et al., 2009).
\*\* Lower and upper bound of the 95% HDI interval of the mean difference between the WEV-model and the model in the row. Negative values indicate that the WEV model performed better.

#### Predicting ERPs from model fits

Based on parameter sets of the WEV-model obtained by fitting the behavioural data, we determined expected ERP amplitudes at the time of P3, ERN, and Pe as a function of SOA and choice accuracy. First, a simple linear transformation was applied to confidence with parameters of the transformation determined based on the EEG data. Fig. 7A shows that the linear transformation of predicted confidence resulted in a reasonably accurate prediction regarding ERP amplitude in the P3 window. In contrast, as can be seen from Fig. 7B, the predicted EEG in the ERN time window was more negative in correct than in incorrect trials, even though the ERN is well-established as a negativity related to errors. The prediction did not match the established polarity of the ERN because the transformation was determined without any assumptions about the polarity of the EEG effects; when the transformation was constraint to a positive relationship between confidence and EEG amplitude, the prediction deviated even more strongly from the observed EEG at the time of the ERN. Likewise, Fig 7C shows that longer SOAs were associated with a positive shift in incorrect trials during the Pe time window, which was just opposite to the pattern observed with confidence judgments (cf. Fig 2B) and therefore was not reproduced by the prediction. As a consequence, there was a medium-sized correction between predicted and observed single-trial amplitudes at the time of the P3, *M* = .33, 95% HDI [.29 .37], while the correlations at the other two time windows were small (ERN: *M* = .09, 95% HDI [.06 .11], Pe: *M* = .16, 95% HDI [.12 .20]). The same results were obtained when we repeated this analysis with the noisy decay model and the detection heuristic model.

The relationship between confidence and ERP amplitudes of course does not need to be linear. For this reason, we fitted non-linear transformations to the data from each subject by assigning the voltage that minimized the prediction error with respect to ERP amplitude to each level of confidence. The only restriction of the transformation was that the relationship between confidence and ERP amplitudes was assumed to be monotonous. Nevertheless, the predictions based on these specifically adapted transformations were only consistent with amplitudes at the time of the P3, but not with ERN or Pe.

## Discussion

The present study revealed a close correlation between decision confidence and EEG activity 350 – 500 after onset of the stimulus, at the time of the P3 component: First, ERP amplitudes at the time of the P3 were directly associated with observers’ confidence judgments. Second, the amplitude at the time of the P3 varied as a function of SOA and choice accuracy in the same way as confidence judgments did. Finally, P3 amplitude could be accurately predicted by the weighted evidence and visibility (WEV) model, which at the same time provided the best account of confidence judgments. In contrast, EEG activity at the time of the ERN component, an established marker of error detection, as well as at the time of the Pe, a marker of error awareness, did not follow the same statistical pattern as decision confidence as a function of SOA and accuracy, despite the fact that a correlation between amplitude and confidence was detected at the time of the Pe. Moreover, there were only weak correlations between the prediction derived from the WEV-model and ERP amplitude at the time of ERN and Pe.

### Bayesian accounts of the relation between P3 and confidence

How can the association between P3 and decision confidence be reconciled with the extended literature on various different roles of the P3? An explanation may be given in terms of the Bayesian brain theory, which states that the brain must use representations about certainty to make optimal perceptual computations (Knill & Pouget, 2004). One possible interpretation in terms of Bayesian Brain theory is that the P3 directly reflects certainty within the decision process (Herding et al., 2019). In line with this interpretation, the P3 showed a statistical pattern theoretically expected to signify confidence in a vibrotactile task (Herding et al., 2019). Moreover, the P3 is related to the accumulation of sensory evidence within the decision process (O’Connell et al., 2012; Twomey et al., 2015). Finally, the P3 is suppressed in highly visible stimuli, even when observers are not required to make a perceptual decision (Pitts, Padwal, Fennelly, Martínez, & Hillyard, 2014). These findings converge with a line of research suggesting that decision confidence may emerge directly from the decision process. For example, neurons in parietal cortex of rhesus monkeys represented both formation of the direction decision and the degree of certainty (Kiani & Shadlen, 2009). Likewise, human EEG correlates of decision formation and confidence coincided in time and in reconstructed sources in a face vs. car discrimination task (Gherman & Philiastides, 2015).

A second interpretation in terms of Bayesian Brain theory is that the P3 reflects sensory representations that include the reliability of the percept (Kopp et al., 2016). This second view is consistent with classical interpretations of the P3 as update of working memory in response to task-relevant events (Donchin & Coles, 1988) or global broadcast of information within a neural global workspace (Sergent et al., 2005). These updated or broadcast representations may encompass the reliability of the percept (Shea & Frith, 2019), which is why the P3 should be correlated with confidence judgments. In line with this interpretation, the WEV-model assumes that confidence is determined by the perceived strength or reliability of the percept based on evidence about choice-relevant and choice-irrelevant features. This means that that the inferred computational principles underlying decision confidence include a representation of the reliability of the percept as well.

### Role of ERN/Pe during the present task?

In the present study, EEG activity in the ERN time window can be interpreted as specifically error detection, but not as decision confidence. EEG activity at the time of the ERN does not reflect confidence because the effects of SOA were opposite to what was expected from observed confidence judgments. At least in the present study, the ERN may not be related to postdecisional sensory evidence, because sensory evidence in correct trials is expected to increase with SOA (Hangya et al., 2016), but at the time of the ERN, the only reliable effect was large negative shift specifically in incorrect trials at the longest SOA. The absence of an ERN at shorter SOAs is in line with a previous study showing that the elicitation of a ERN requires participants to know which response is the correct one (Di Gregorio et al., 2018). Likewise, in the present study, observers also did not know for sure which response had been correct at shorter SOAs because the mask impeded perception of the target. These findings are also consistent with a previous study showing that the ERN occur only when observers make erroneous responses to stimuli rated as “visible” (Charles et al., 2014, 2013). Although we did not measure conscious awareness in the present study, we can extrapolate from other studies using the same task that observers’ conscious percepts of the stimuli were degraded in shorter SOAs (Rausch & Zehetleitner, 2019a; Zehetleitner & Rausch, 2013); possibly, weakly conscious stimuli are not sufficient to trigger an ERN.

A possible interpretation for the role of the Pe in the present study is as accumulation of postdecisional sensory evidence. At least in the present study, the Pe does not reflect decision confidence because their statistical patterns as functions of SOA and choice accuracy are not compatible. In addition, the Pe does not exclusively reflect error awareness, because EEG activity at the time of the Pe was correlated with confidence in correct trials. However, the pattern of the Pe as a function of SOA and choice accuracy matches the diverging pattern between correct and incorrect responses expected from postdecisional accumulation of sensory evidence (Moran et al., 2015). The contribution of postdecisional sensory evidence to confidence may be relatively small in the present paradigm, because the mask prevents ongoing accumulation of evidence from sensory memory. Accordingly, cognitive modelling showed that the WEV model fitted confidence much better than the SDT model with postdecisional evidence. If the Pe reflects postdecisional accumulation of evidence, this explains why in the present study effects at the time of the Pe seemed to be limited to high confidence trials. The efficiency of the mask varies across trials, and presumably the mask had been relatively ineffective in trials when observers reported high degrees of confidence. Moreover, if the Pe represents postdecisional sensory evidence, it can be explained why a previous study detected an association between the Pe and all degrees of confidence (Boldt & Yeung, 2015). As stimuli in that study were not masked, postdecisional accumulation of sensory evidence may have been more effective than in the present study. Finally, the Pe may not only be sensitive to postdecisional sensory evidence, but may reflect also other sources of information, including response conflict, efference copy, proprioception, perception of action effects, and interoception (Ullsperger et al., 2010; Wessel et al., 2011).

### Statistical signatures of confidence?

The present study demonstrates that statistical patterns of confidence can provide a strong test for identifying correlates of confidence, although it is crucial to validate statistical signatures of confidence empirically by behavioural measures of confidence. It has been argued that if confidence is determined objectively as the posterior probability of being correct, the pattern referred to as folded X-pattern is the statistical signature of confidence (Hangya et al., 2016; Sanders et al., 2016). Therefore, a substantial number of recent studies have searched for the folded X-pattern to empirically identify correlates of decision confidence (Braun et al., 2018; Fetsch et al., 2014; Herding et al., 2019; Lak et al., 2017; Sanders et al., 2016; Urai et al., 2017). However, it has been shown mathematically that the folded X-pattern is neither a necessary nor a sufficient condition for Bayesian confidence (Adler & Ma, 2018; Rausch & Zehetleitner, 2019b). The present study showed empirically that a second statistical pattern of confidence exists and can be used to identify correlates of confidence. Had we not measured decision confidence directly and relied on the purported folded-X signature, the Pe, not the P3, would have been falsely considered a correlate of confidence.

## Conclusion

The present results suggest that there is no single EEG correlate of decision confidence and error awareness. EEG activity over parietal electrodes 350 – 500 ms after onset of the stimulus is closely correlated with decision confidence. However, EEG components after the response, which have been established as markers of error detection or error awareness, are dissociated from decision confidence.

## Acknowledgements

We are grateful to Sebastian Hellmann for help in programming the experiment, to Leonie Manzke for help with Python, and Marina Nickl, Anna Bogner, and Johanna Dietrich for help with data collection.

## References

Adler, W. T., & Ma, W. J. (2018). Limitations of proposed signatures of Bayesian confidence. Neural Computation, 30(12), 3327–3354. https://doi.org/10.1162/neco_a_01141

Aitchison, L., Bang, D., Bahrami, B., & Latham, P. E. (2015). Doubly Bayesian Analysis of Confidence in Perceptual Decision-Making. PLoS Computational Biology, 11(10), 1–23. https://doi.org/10.1371/journal.pcbi.1004519

Akaike, H. (1974). A New Look at the Statistical Model Identification. IEEE Transactions on Automatic Control, AC-19(6), 716–723. https://doi.org/10.1007/978-1-4612-1694-0_16

Barrett, A. B., Dienes, Z., & Seth, A. K. (2013). Measures of metacognition on signal-detection theoretic models. Psychological Methods, 18(4), 535–552. https://doi.org/10.1037/a0033268

Boldt, A., & Yeung, N. (2015). Shared Neural Markers of Decision Confidence and Error Detection. The Journal of Neuroscience, 35(8), 3478–3484. https://doi.org/10.1523/JNEUROSCI.0797-14.2015

Braun, A., Urai, A. E., & Donner, T. H. (2018). Adaptive History Biases Result from Confidence-weighted Accumulation of Past Choices. The Journal of Neuroscience, 38(10), 2418–2429. https://doi.org/10.1523/JNEUROSCI.2189-17.2017

Burnham, K. P., & Anderson, D. R. (2002). Model selection and multimodel inference: A practical information--theoretic approach (2nd ed.). New York: Springer.

Charles, L., King, J.-R., & Dehaene, S. (2014). Decoding the Dynamics of Action, Intention, and Error Detection for Conscious and Subliminal Stimuli. The Journal of Neuroscience, 34(4), 1158–1170. https://doi.org/10.1523/JNEUROSCI.2465-13.2014

Charles, L., Opstal, F. Van, Marti, S., & Dehaene, S. (2013). Distinct brain mechanisms for conscious versus subliminal error detection. NeuroImage, 73, 80–94.

Charles, L., & Yeung, N. (2018). Dynamic sources of evidence supporting confidence judgments and error detection. Journal of Experimental Psychology: Human Perception and Performance, 45(1), 39–52.

Di Gregorio, F., Maier, M. E., & Steinhauser, M. (2018). Errors can elicit an error positivity in the absence of an error negativity: Evidence for independent systems of human error monitoring. NeuroImage, 172, 427–436. https://doi.org/10.1016/j.neuroimage.2018.01.081

Donchin, E., & Coles, M. G. (1988). Is the P300 component a manifestation of context updating? Behavioral and Brain Sciences, 11(3), 357–374.

Eimer, M., & Mazza, V. (2005). Electrophysiological correlates of change detection. Psychophysiology, 42(3), 328–342.

Falkenstein, M., Hohnsbein, J., Hoormann, J., & Blanke, L. (1991). Effects of crossmodal divided attention on ERP components: Error processing in choice reaction tasks. Electroencephalography and Clinical Neurophysiology, 78(6), 447–455. https://doi.org/10.1016/0013-4694(91)90061-8

Fetsch, C. R., Kiani, R., Newsome, W. T., & Shadlen, M. N. (2014). Effects of Cortical Microstimulation on Confidence in a Perceptual Decision. Neuron, 83(4), 797–804. https://doi.org/10.1016/j.neuron.2014.07.011

Gehring, W. J., Goss, B., Coles, M. G. H., Meyer, D. E., & Donchin, E. (1993). A neural system for error detection and compensation. Psychological Science, 4(6), 385–390.

Gherman, S., & Philiastides, M. G. (2015). Neural representations of confidence emerge from the process of decision formation during perceptual choices. NeuroImage, 106, 134–143. https://doi.org/10.1016/j.neuroimage.2014.11.036

Gramfort, A., Luessi, M., Larson, E., Engemann, D. A., Strohmeier, D., Brodbeck, C., … Hämäläinen, M. (2013). MEG and EEG data analysis with MNE-Python. Frontiers in Neuroscience, 7(267), 1–13. https://doi.org/10.3389/fnins.2013.00267

Gramfort, A., Luessi, M., Larson, E., Engemann, D., Strohmeier, D., Brodbeck, C., … Hämäläinen, M. (2014). MNE software for processing MEG and EEG data. NeuroImage, 86, 446–460. https://doi.org/10.1021/nl061786n.Core-Shell

Green, D. M., & Swets, J. A. (1966). Signal detection theory and psychophysics. New York: Wiley.

Hangya, B., Sanders, J. I., & Kepecs, A. (2016). A Mathematical Framework for Statistical Decision Confidence. Neural Computation, 28(9), 1840–1858. https://doi.org/10.1162/NECO_a_00864

Herding, J., Ludwig, S., von Lautz, A., Spitzer, B., & Blankenburg, F. (2019). Centro-parietal EEG potentials index subjective evidence and confidence during perceptual decision making. NeuroImage, 201, 116011. https://doi.org/10.1016/j.neuroimage.2019.116011

Hillyard, S. A., Squires, K. C., Bauer, J. W., & Lindsay, P. H. (1971). Evoked Potential Correlates of Auditory Signal Detection. Science, 172, 1357–1360.

Hyvärinen, A. (1999). Fast and robust fixed-point algorithms for independent component analysis. IEEE Transactions on Neural Networks, 10(3), 626–634. https://doi.org/10.1109/72.761722

Kellen, D., & Klauer, K. C. (2015). Signal Detection and Threshold Modeling of Confidence-Rating ROCs: A Critical Test With Minimal Assumptions. Psychological Review, 122(3), 542–557.

Kepecs, A., Uchida, N., Zariwala, H., & Mainen, Z. F. (2008). Neural correlates, computation and behavioural impact of decision confidence. Nature, 455(7210), 227–231. https://doi.org/10.1038/nature07200

Kiani, R., Corthell, L., & Shadlen, M. N. (2014). Choice Certainty Is Informed by Both Evidence and Decision Time. Neuron, 84(6), 1329–1342. https://doi.org/10.1016/j.neuron.2014.12.015

Kiani, R., & Shadlen, M. N. (2009). Representation of confidence associated with a decision by neurons in the parietal cortex. Science, 324(5928), 759–764. https://doi.org/10.1126/science.1169405

Knill, D. C., & Pouget, A. (2004). The Bayesian brain: The role of uncertainty in neural coding and computation. Trends in Neurosciences, 27(12), 712–719. https://doi.org/10.1016/j.tins.2004.10.007

Koivisto, M., & Revonsuo, A. (2010). Event-related brain potential correlates of visual awareness. Neuroscience and Biobehavioral Reviews, 34, 922–934. https://doi.org/10.1016/j.neubiorev.2009.12.002

Kopp, B., Seer, C., Lange, F., Kluytmans, A., Kolossa, A., Fingscheidt, T., & Hoijtink, H. (2016). P300 amplitude variations, prior probabilities, and likelihoods: A Bayesian ERP study. Cognitive, Affective and Behavioral Neuroscience, 16(5), 911–928. https://doi.org/10.3758/s13415-016-0442-3

Lak, A., Nomoto, K., Keramati, M., Sakagami, M., & Kepecs, A. (2017). Midbrain Dopamine Neurons Signal Belief in Choice Accuracy during a Perceptual Decision. Current Biology, 27(6), 821–832. https://doi.org/10.1016/j.cub.2017.02.026

Lamy, D., Salti, M., & Bar-haim, Y. (2008). Neural Correlates of Subjective Awareness and Unconscious Processing: An ERP Study. Journal of Cognitive Neuroscience, 21(7), 1435–1446.

Lee, M. D., & Wagenmakers, E.-J. (2013). Bayesian Cognitive Modeling: A Practical Course. Cambridge, UK: Cambridge University Press.

Luck, S. J. (2014). Comparing Conditions with Different Numbers of Trials or Different Levels of Noise. In An Introduction to the Event-Related Potential Technique. MIT Press. Retrieved from http://mitp-content-server.mit.edu:18180/books/content/sectbyfn?collid=books_pres_0&id=8575&fn=8575_009a_0.pdf

Macmillan, N. A., & Creelman, C. D. (2005). Detection Theory. A user’s guide. Mahwah, NY: Lawrence Erlbaum Associates.

Maniscalco, B., & Lau, H. (2016). The signal processing architecture underlying subjective reports of sensory awareness. Neuroscience of Consciousness, 1, niw002. https://doi.org/10.1093/nc/niw002

Maniscalco, B., Peters, M. A. K., & Lau, H. (2016). Heuristic use of perceptual evidence leads to dissociation between performance and metacognitive sensitivity. Attention, Perception & Psychophysics, 78, 923–937. https://doi.org/10.3758/s13414-016-1059-x

Moran, R., Teodorescu, A. R., & Usher, M. (2015). Post choice information integration as a causal determinant of confidence: Novel data and a computational account. Cognitive Psychology, 78, 99–147. https://doi.org/10.1016/j.cogpsych.2015.01.002

Morey, R. D., & Rouder, J. N. (2015). BayesFactor: Computation of Bayes factors for common designs. R package version 0.9.10-1. Retrieved from https://cran.r-project.org/package=BayesFactor

Nelder, J. A., & Mead, R. (1965). A simplex method for function minimization. The Computer Journal, 7, 308--313.

Nieuwenhuis, S., de Geus, E. J., & Aston-Jones, G. (2011). The anatomical and functional relationship between the P3 and autonomic components of the orienting response. Psychophysiology, 48(2), 162–175. https://doi.org/10.1111/j.1469-8986.2010.01057.x.The

Nieuwenhuis, S., Ridderinkhof, R. K., Blom, J., Band, G. P. H., & Kok, A. (2001). Error-related brain potentials are differentially related to awareness of response errors: Evidence from an antisaccade task. Psychophysiology, 38(5), 752–760. https://doi.org/10.1017/S0048577201001111

O’Connell, R. G., Dockree, P. M., & Kelly, S. P. (2012). A supramodal accumulation-to-bound signal that determines perceptual decisions in humans. Nature Neuroscience, 15(12), 1729–1735. https://doi.org/10.1038/nn.3248

Overbeek, T. J. M., Nieuwenhuis, S., & Ridderinkhof, K. R. (2005). Dissociable components of error processing: On the functional significance of the Pe vis-à-vis the ERN/Ne. Journal of Psychophysiology, 19(4), 319–329. https://doi.org/10.1027/0269-8803.19.4.319

Peirce, J. W. (2007). PsychoPy: Psychophysics software in Python. Journal of Neuroscience Methods, 162, 8–13. https://doi.org/10.1016/j.jneumeth.2006.11.017

Peirce, J. W. (2009). Generating stimuli for neuroscience using PsychoPy. Frontiers in Neuroinformatics, 2(10), 1–8. https://doi.org/10.3389/neuro.11.010.2008

Perrin, F., Pernier, J., Bertrand, O., & Echallier, J.. (1989). Spherical splines for scalp potential and current density mapping. Electroencephalography and Clinical Neurophysiology, 72(6), 184–187.

Peters, M. A. K., Thesen, T., Ko, Y. D., Maniscalco, B., Carlson, C., Davidson, M., … Lau, H. (2017). Perceptual confidence neglects decision-incongruent evidence in the brain. Nature Human Behaviour, 1(7), 1–21. https://doi.org/10.1038/s41562-017-0139

Philiastides, M. G., Heekeren, H. R., & Sajda, P. (2014). Human scalp potentials reflect a mixture of decision-related signals during perceptual choices. Journal of Neuroscience, 34(50), 16877–16889. https://doi.org/10.1523/JNEUROSCI.3012-14.2014

Pitts, M. A., Padwal, J., Fennelly, D., Martínez, A., & Hillyard, S. A. (2014). Gamma band activity and the P3 reflect post-perceptual processes, not visual awareness. NeuroImage, 101, 337–350. https://doi.org/10.1016/j.neuroimage.2014.07.024

Pleskac, T. J., & Busemeyer, J. R. (2010). Two-Stage Dynamic Signal Detection: A Theory of Choice, Decision Time, and Confidence. Psychological Review, 117(3), 864–901. https://doi.org/10.1037/a0019737

Pouget, A., Drugowitsch, J., & Kepecs, A. (2016). Confidence and certainty: distinct probabilistic quantities for different goals. Nature Neuroscience, 19(3), 366–374. https://doi.org/10.1038/nn.4240

Rausch, M., Hellmann, S., & Zehetleitner, M. (2018). Confidence in masked orientation judgments is informed by both evidence and visibility. Attention, Perception, & Psychophysics, 80(1), 134–154. https://doi.org/10.3758/s13414-017-1431-5

Rausch, M., & Zehetleitner, M. (2017). Should metacognition be measured by logistic regression? Consciousness and Cognition, 49, 291–312. https://doi.org/10.1016/j.concog.2017.02.007

Rausch, M., & Zehetleitner, M. (2019a). Modelling visibility judgments using models of decision confidence. PsyArXiv. https://doi.org/10.31219/osf.io/7dakz

Rausch, M., & Zehetleitner, M. (2019b). The folded X-pattern is not necessarily a statistical signature of decision confidence. PLoS Computational Biology, 15(10), e1007456. https://doi.org/10.1371/journal.pcbi.1007456

Resulaj, A., Kiani, R., Wolpert, D. M., & Shadlen, M. N. (2009). Changes of mind in decision-making. Nature, 461, 263–266. https://doi.org/10.1038/nature08275

Rolls, E. T., Grabenhorst, F., & Deco, G. (2010). Decision-Making, Errors, and Confidence in the Brain. Journal of Neurophysiology, 104(5), 2359–2374. https://doi.org/10.1152/jn.00571.2010

Rouder, J. N., & Morey, R. D. (2012). Default Bayes Factors for Model Selection in Regression. Multivariate Behavioral Research, 47(6), 877–903.

Rouder, J. N., Speckman, P. L., Son, D., & Morey, R. D. (2009). Bayesian t tests for accepting and rejecting the null hypothesis. Psychonomic Bulletin & Review, 16(2), 225–237. https://doi.org/10.3758/PBR.16.2.225

Sanders, J. I., Hangya, B., & Kepecs, A. (2016). Signatures of a Statistical Computation in the Human Sense of Confidence. Neuron, 90, 499–506. https://doi.org/10.1016/j.neuron.2016.03.025

Scheffers, M. K., & Coles, M. G. H. (2000). Performance monitoring in a confusing world: Error-related brain activity, judgments of response accuracy, and types of errors. Journal of Experimental Psychology: Human Perception and Performance, 26(1), 141–151. https://doi.org/10.1037/0096-1523.26.1.141

Schwarz, G. (1978). Estimating the dimensions of a model. The Annals of Statistics, 6(2), 461–464. https://doi.org/10.1214/aos/1176348654

Sergent, C., Baillet, S., & Dehaene, S. (2005). Timing of the brain events underlying access to consciousness during the attentional blink. Nature Neuroscience, 8(10), 1391–1400. https://doi.org/10.1038/nn1549

Shea, N., & Frith, C. D. (2019). The Global Workspace Needs Metacognition. Trends in Cognitive Sciences, 23(7), 560–571. https://doi.org/10.1016/j.tics.2019.04.007

Steinhauser, M., Maier, M., & Hübner, R. (2008). Modeling Behavioral Measures of Error Detection in Choice Tasks: Response Monitoring Versus Conflict Monitoring. Journal of Experimental Psychology: Human Perception and Performance, 34(1), 158–176.

Steinhauser, M., & Yeung, N. (2010). Decision Processes in Human Performance Monitoring. The Journal of Neuroscience, 30(46), 15643–15653. https://doi.org/10.1523/JNEUROSCI.1899-10.2010

Steinhauser, M., & Yeung, N. (2012). Error awareness as evidence accumulation: Effects of speed-accuracy trade-off on error signaling. Frontiers in Human Neuroscience, 6(240), 1–12. https://doi.org/10.3389/fnhum.2012.00240

Stolyarova, A., Rakhshan, M., Hart, E. E., O’Dell, T. J., Peters, M. A. K., Lau, H., … Izquierdo, A. (2019). Dissociable roles for Anterior Cingulate Cortex and Basolateral Amygdala in Decision Confidence and Learning under Uncertainty. Nature Communications, 10(4704), 1–14. https://doi.org/10.1101/655860

Tagliabue, C. F., Veniero, D., Benwell, C. S. Y., Cecere, R., Savazzi, S., & Thut, G. (2019). Subjective perceptual experience tracks the neural signature of sensory evidence accumulation during decision formation. Scientific Reports, 9(4949), 1–12. https://doi.org/10.1038/s41598-019-41024-4

Twomey, D. M., Murphy, P. R., Kelly, S. P., & O’Connell, R. G. (2015). The classic P300 encodes a build-to-threshold decision variable. European Journal of Neuroscience, 42(1), 1636–1643. https://doi.org/10.1111/ejn.12936

Ullsperger, M., Harsay, H. A., Wessel, J. R., & Ridderinkhof, K. R. (2010). Conscious perception of errors and its relation to the anterior insula. Brain Structure & Function, 214(5–6), 629–643. https://doi.org/10.1007/s00429-010-0261-1

Urai, A. E., Braun, A., & Donner, T. H. (2017). Pupil-linked arousal is driven by decision uncertainty and alters serial choice bias. Nature Communications, 8, 14637. https://doi.org/10.1038/ncomms14637

van den Berg, R., Anandalingam, K., Zylberberg, A., Kiani, R., Shadlen, M. N., & Wolpert, D. M. (2016). A common mechanism underlies changes of mind about decisions and confidence. ELife, 5, e12192. https://doi.org/10.7554/eLife.12192

Verleger, R., Ja, P., & Wascher, E. (2005). Evidence for an Integrative Role of P3b in Linking Reaction to Perception. Journal of Psychophysiology, 20(2), 1–17. https://doi.org/10.1027/0269-8803.19.2.xxx

Vidal, F., Burle, B., Bonnet, M., Grapperon, J., & Hasbroucq, T. (2003). Error negativity on correct trials: a reexamination of available data. Biological Psychology, 64, 265–282. https://doi.org/10.1016/S0301-0511(03)00097-8

Wessel, J. R., Danielmeier, C., & Ullsperger, M. (2011). Error Awareness Revisited: Accumulation of Multimodal Evidence from Central and Autonomic Nervous Systems. Journal of Cognitive Neuroscience, 23(10), 3021–3036. https://doi.org/10.1162/jocn.2011.21635

Wickens, T. D. (2002). Elementary signal detection theory. New York: Oxford University Press.

Zehetleitner, M., & Rausch, M. (2013). Being confident without seeing: What subjective measures of visual consciousness are about. Attention, Perception, and Psychophysics, 75(7), 1406–1426. https://doi.org/10.3758/s13414-013-0505-2

